# Studying relative RNA localization From nucleus to the cytosol

**DOI:** 10.1101/2024.03.06.583744

**Authors:** Vasilis F. Ntasis, Roderic Guigó

## Abstract

The precise coordination of important biological processes, such as differentiation and development, is highly dependent on the regulation of expression of the genetic information. The flow of the genetic information is tightly regulated on multiple levels. Among them, RNA export to cytosol is an essential step for the production of proteins in eukaryotic cells. Hence, estimating the relative concentration of RNA molecules of a given transcript species in the nucleus and in the cytosol is of major significance as it contributes to the understanding of the dynamics of RNA trafficking between the nucleus and the cytosol. The most efficient way to estimate the levels of RNA species genome-wide is through RNA sequencing (RNAseq). While RNAseq can be performed separately in the nucleus and in the cytosol, because measured transcript levels are relative to the total volume of RNA in these compartments, and because this volume is usually unknown, the transcript levels in the nucleus and in the cytosol cannot be directly compared. Here we show theoretically that if, in addition to nuclear and cytosolic RNA-seq, whole cell RNA-seq is also performed, then accurate estimations of the localization of transcripts can be obtained. Based on this, we designed a method that estimates, first the fraction of the total RNA volume in the cytosol (nucleus), and then, this fraction for every transcript. We evaluate our methodology on simulated data and nuclear and cytosolic single cell data available. Finally, we use our method to investigate the cellular localization of transcripts using bulk RNAseq data from the ENCODE project.

## Introduction

Estimating RNA subcellular localization is crucial for comprehending different cellular processes and their biological implications. For instance, the regulation of the amount of protein produced can be achieved through nuclear retention of RNA molecules. By localizing certain transcripts in the nucleus, they can be prevented from being translated into proteins, thereby controlling protein abundance and ensuring the precision of regulatory mechanisms (Mauger et al. 2016; Naro et al. 2017). RNA localization can also buffer protein levels from bursty transcription, and protect from transcripts with retain introns, as it enables a more balanced and controlled protein production, minimizing fluctuations and maintaining cellular homeostasis (Lécuyer et al. 2007; Bahar Halpern et al. 2015; Battich et al. 2015). Other processes in which RNA subcellular localization plays an important role include transcription regulation (Syed et al. 2021), nuclear size control (Kume et al. 2017; Cantwell and Nurse 2019), and protein localization (Lécuyer et al. 2007; Martin and Ephrussi 2009). Finally, the disruption of RNA localization pathways has been implicated in different diseases, particularly in various neuromuscular disorders (Cooper et al. 2009). So, determining RNA localization is essential for understanding how cells work.

The most crucial subcellular RNA segregation in eukaryotes is the one between the nucleus and cytosol. Different approaches have been employed to infer the preferential nuclear vs cytosolic localization of transcripts. Likely the earliest and most widely used methods, such as “in situ” hybridization, rely on oligonucleotide hybridization coupled to imaging (see Carlevaro-Fita and R Johnson 2019, for a review). These methods, however, are low throughput, and can monitor simultaneously only up to hundreds of transcripts. Furthermore, efforts have been made to develop computational methods for predicting RNA subcellular localization (see J Wang et al. 2023, for a review). Nevertheless, most of these models base their predictions in sequence features, and are limited by the data they have been trained on. Hence, they may miss context specific changes in RNA localization that are independent of transcript sequence.

RNA sequencing (RNAseq), on the other hand, allows for monitoring the abundance of all (or most) transcripts within an RNA sample. Recent advancements have introduced innovative methods that combine proximity labeling with RNAseq (Fazal et al. 2019; Padrón et al. 2019; P Wang et al. 2019). These approaches utilize engineered proteins that label molecules in proximity, facilitating the identification of interactions and spatial associations between nuclear and cytosolic components. They also have their own limitations. For instance, they are unsuitable for studying human tissues, because of their dependence on the use of recombinant proteins, or for studying RNAs sequestered within large complexes, due to their steric inaccessibility.

If RNAseq is performed, following cellular fractionation (i.e. the isolation of the nuclear and cytosolic RNA fractions), it could, in principle, produce separate estimates of transcript abundances in the nucleus and in the cytosol (Djebali et al. 2012; Lefebvre et al. 2017). Measurements of transcript abundances from typical RNAseq experiments, however, are relative to the total amount of RNA in the biological sample sequenced. Since the sequencing depth in these experiments is usually independent of the total amount of RNA, estimates of transcript abundances (FPKMs, RPKMs, TPMs, etc.) cannot be directly related to the real counts of RNA molecules of the different transcript species. For instance, if two different RNA samples, containing 103 and 106 RNA molecules respectively are both sequenced to the same number of reads, a given transcript capturing 1% of the total RNA in both samples, will be estimated to have exactly the same expression in both samples, even though it will be in ten copies in the first and in 10,000 copies in the second.

Strictly speaking, therefore, RNAseq estimates of transcript abundances cannot be compared across samples. In practice, however, for most differential gene expression analysis this has little impact, as usually the amount of RNA of the compared biological conditions is roughly the same (as when compared normal vs diseased conditions), and, from the biological standpoint, relative abundances of transcripts may capture more appropriately the stoichiometry of biochemical reactions than absolute values, and changes in these abundances, therefore, may more strongly correlate with associated phenotypic changes. This is not the case, however, when comparing RNAseq based estimates of transcript abundances from different cellular fractions (i.e. nucleus vs cytosol). In this case, as the total RNA volume may radically differ across fractions, relative estimates within each fraction provide little information of the differences in abundances of a given transcript in different fractions, and therefore, these estimates cannot be used to study the subcellular (i.e. nucleo-cytoplasmic) distribution of transcripts.

To address these limitations, Carlevaro-Fita and R Johnson 2019 sketched a method to estimate absolute RNA localization from fractionated RNAseq. The method models total RNA expression as the sum of nuclear and cytoplasmic concentrations. Here, we further explore these ideas, and we first show that from matched nuclear, cytosolic and whole cell RNAseq data it is possible to estimate the fraction of total cellular RNA that localizes to the cytosol (or to the nucleus). We show next that from this fraction and from relative measurements of transcript abundances (i.e. FPKMs), it is possible to estimate the nuclear vs cytosolic ratio of the number of molecules of a given transcript. To obtain these estimates, we developed a method that employs Bayesian linear regression. We benchmark this method using simulated data and recently produced single cell data, and we show that it provides good estimates of the nucleo-cytosolic molecular ratios of transcripts. We use the method to investigate the nucleo-cytoplasmic distribution of transcripts in a number of ENCODE cell lines for which whole cell, nuclear and cytosolic RNAseq are all available. We report that this distribution is a characteristic of different transcript types, and that it is partially associated with sequence features of transcripts. Consistently, we found that subcellular localization is a property of the transcripts rather than of the genes, and that many genes have transcripts that are consistently localized in different cellular compartments. We also found that, contrary to the expectation from purely statistical considerations, constitutively expressed transcripts tend to have a more consistent cellular localization across biological conditions than transcripts expressed in a more cell-type specific manner.

## Results

### The transcript localization index

Given a eukaryotic cell (or a population of cells), let *m*_*f*_ (*i*) denote the number of RNA molecules of transcript *i* (*i* = 1, 2, 3, …, *t*) in cell fraction *f*, where here *f ∈ {w, n, c}*, referring to whole-cell (*w*), nuclear (*n*), and cytosolic (*c*) fraction respectively. Then, the total number of molecules of a given transcript *i* in the whole-cell is equal to the sum of the number of molecules of that transcript in the nucleus and in the cytosol:

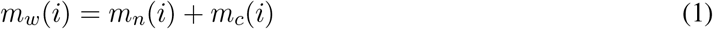

Here, our aim is to estimate the proportion of the total RNA molecules of a given transcript localized in the nucleus, or in the cytosol:

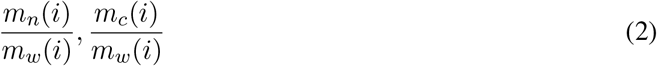

We will show first how these proportions can be estimated using whole cell together with fractional RNAseq data. Let’s assume that we perform a RNAseq experiment in fraction f, with a total of *D*_*f*_ sequenced reads. The expected number of reads mapping to transcript *i, r*_*f*_ (*i*), is proportional to the number of molecules and the length of transcript *i*. That is:

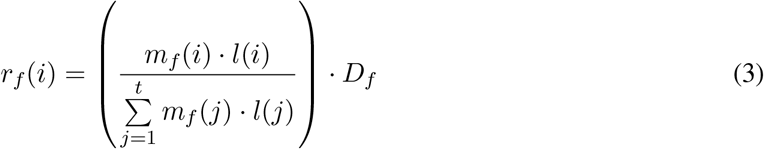

where *l*(*i*) is the length of transcript *i*. Given equation (3), equation (1) can be re-written as:

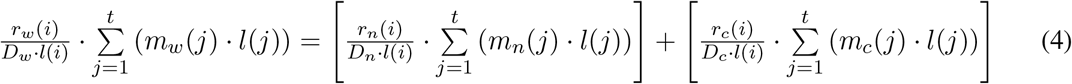

which is equivalent to:

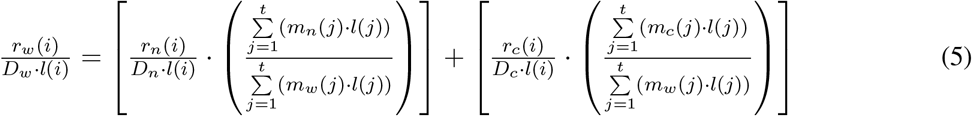

Here, the 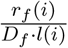 quantities are the normalized read count for transcript *i*. A scaled version of that is the widely used *FPKM* quantity (fragments per kilobase of transcript per million fragments mapped in a pairedend sequencing experiment). Hence, equation (5) can be rewritten as follows:

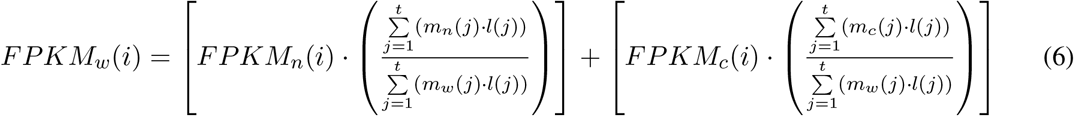

The 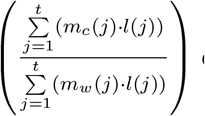 quantity is the fraction of the total cellular RNA volume (i.e. the total number of nucleotides in RNA molecules) localized in the cytosol, and 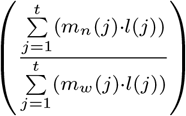 is the fraction localized in the nucleus. Therefore, if *β* denotes the cytosolic fraction, equation (6) can be written as:

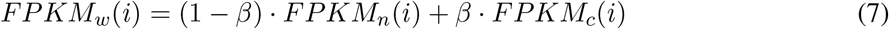

Finally, if *β* is known, it can then be used to calculate the fraction of the total RNA molecules of a specific transcript *i* that are localized in the cytosol (eq. 2) from its cytosolic and nuclear FPKM values:

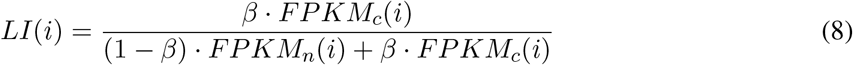

We call this the “localization index” of a given transcript. When *β* = 0.5, the localization index simplifies to:

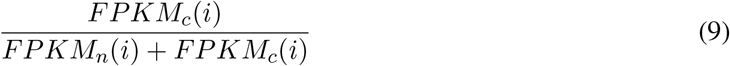

We refer to this ratio as the “naive localization index”. Often, the naive localization index, or similarly the 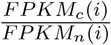 ratio, has been used to quantify transcript localization. However, as we will show next, as *β* deviates from 0.5, these ratios (based exclusively on FPKM values), under(*β >* 0.5) or overestimate (*β <* 0.5) the true localization index.

### Conceptual example

We made a thought experiment to illustrate the rationale above in Figure 1. We assume two scenarios corresponding to two different cell types. In the first, the amount of RNA in the sample to be sequenced is the same in the nucleus and in the cytosol (*β* = 0.5). In the second, 80% of the RNA is in the cytosol (*β* = 0.8, Fig. 1A). We assume that sequencing is performed independently in the nucleus and in the cytosol, and read counts are assigned to genes proportionally to the genes’ molecular counts as in equation (3) (Fig. 1B).

**Figure 1.**
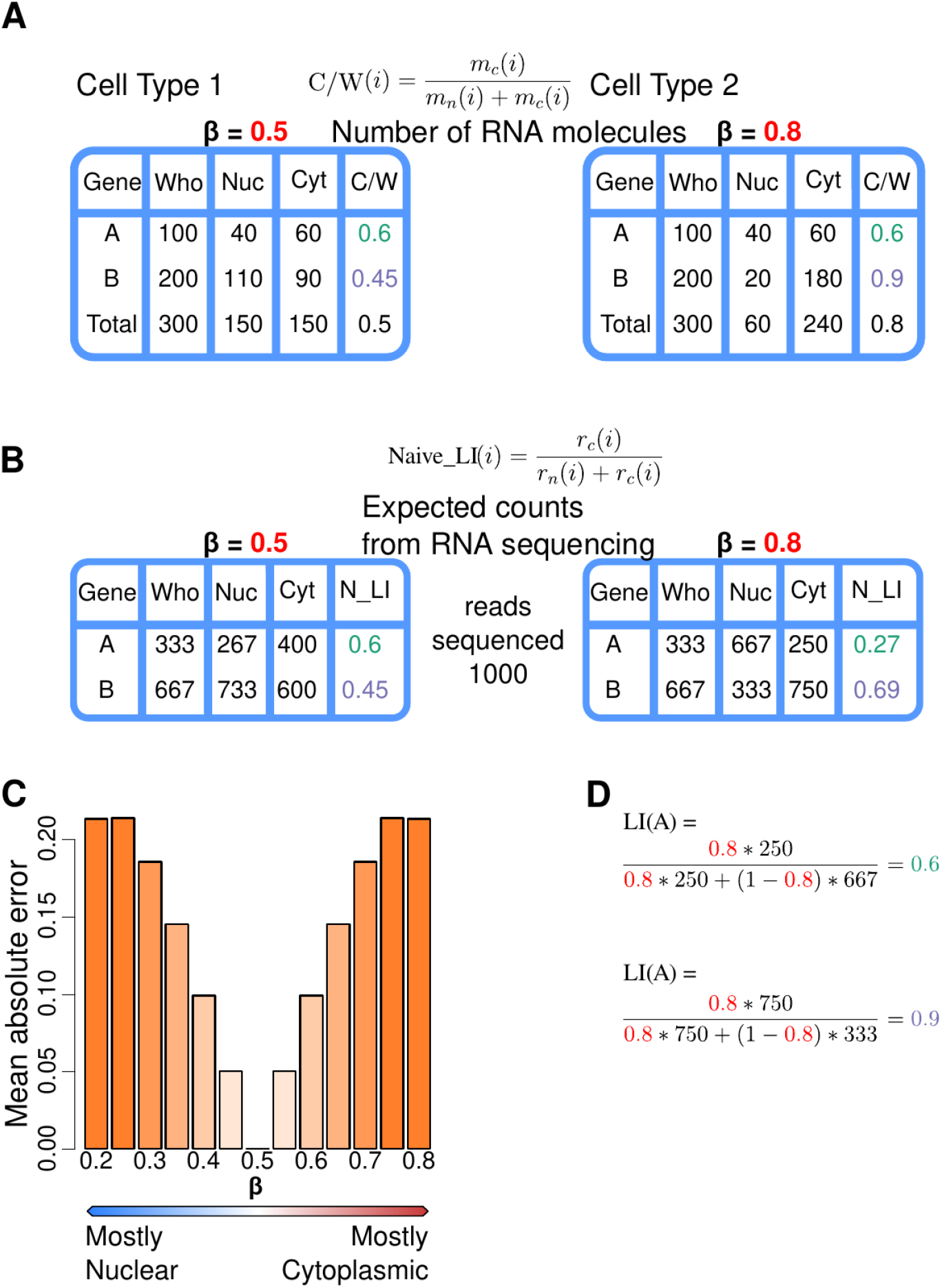
Let’s assume a genome made of two genes A and B of the same transcript length. (A) Let’s assume that we measure their expression in two cell types (1 and 2). In both cell types, whole cells contain 100 molecules of gene A and 200 molecules of gene B. Gene A has identical nucleo-cytosolic molecular distribution in the two cell types (40 molecules in the nucleus and 60 in the cytosol), while gene B is preferentially nuclear in cell type 1 (110 vs 90 molecules), and preferentially cytosolic in cell type 2 (20 vs 180 molecules). The proportion of the total RNA molecules of gene A localized in the cytosol is 0.6 in both cell types, while the corresponding proportion for gene B is 0.45 in cell type 1 and 0.9 in cell type 2. Thus, in cell type 1 the amount of RNA is the same in the nucleus than in the cytosol (*β* = 0.5), while in cell type 2, 80% of the total RNA is in the cytosol (*β* = 0.8). (B) Let’s assume that we perform nuclear and cytosolic RNAseq experiments in each cell type at a sequence depth of 1,000 reads each experiment. We expect the reads to be distributed proportionally to the molecular counts as in equation (3). Based on the nuclear and cytosolic read counts, we can calculate the naive localization index (eq. 9), here expressed in terms of read counts since the sequencing depth is constant across all the samples sequenced. In cell type 1 (with nucleo-cytosolic equidistribution of RNA), the naive localization index (*N* _*LI*) is identical to the portion of the total RNA molecules of each gene localized in the cytosol (*C/W*). In cell type 2, in contrast, where 80% of the cellular RNA is in the cytosol, the naive localization index (0.27 for gene A and 0.69 for gene B) deviates substantially from the true proportions. Actually, the naive localization index for gene A suggests that the gene is preferentially nuclear in cell type 2, even though it occurs in a larger number of copies in the cytosol. (C) The deviation of the naive localization index from the actual portion of the total RNA molecules of a specific gene localized in the cytosol increases as the fraction of the total RNA volume localized in the cytosol deviates from *β* = 0.5. In the same setup of a hypothetical genome of two genes, we measured this deviation for varying values of *β*. (D) If, in addition of the nuclear and cytosolic RNAseq experiments, we perform a whole cell RNAseq experiment (for instance, at the same depth of 1,000 reads), the derived read counts for genes A and B can be used to estimate *β*. This, in turn, can be employed to calculate the localization index (LI, eq. 8) of the genes in cell type 2, which is identical to the portion of the total RNA molecules of a gene localized in the cytosol (*C/W*).

In the first cell type (*β* = 0.5), the naive localization index (eq. 9) produces the correct estimations of the portion of the total RNA molecules of a given transcript localized in the cytosol (eq. 2). In cell type 2, however, these estimates differ substantially from the actual values, and they may even suggest the wrong preferential cellular localization. This deviation increases as the fraction of the total RNA volume localized in the cytosol (or nucleus) diverges from *β* = 0.5 (Fig. 1C).

If, in addition to the nuclear and cytosolic RNAseq experiments, we perform a whole cell RNAseq experiment, then we can estimate *β* (as we show in the next section). This, in turn, can be employed to compute the localization index (LI), which in all cases is identical to the portion of the total RNA molecules of a given transcript localized in the cytosol (Fig. 1D).

### Estimation of the *β*

In most experimental setups *β*, the fraction of the total RNA volume localized in the cytosol, is unknown. However, as indicated above (eq. 7), it is possible to estimate it, if there are whole cell, nuclear, and cytosolic RNAseq data available. Ideally, *β* is the same for all transcripts. However, because of measurement errors intrinsic to RNAseq experiments, this is unlikely to be the case. Thus, we formulated the relationship in equation (7) as a regression problem:

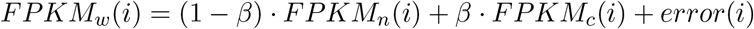

The regression was implemented under a Bayesian framework, and the error term was modeled as a Student’s t-distribution, so as to generate more robust estimates. We utilized the Stan probabilistic programming language (Stan Development Team 2021), in order to employ our Bayesian linear regression model, and combined it with Nextflow workflow manager (Tommaso et al. 2017). We created Stan-NF (https://github.com/vntasis/stan-nf), a Nextflow pipeline that uses CmdStan to compile a Stan model, run MCMC sampling, and compute diagnostics, in a reproducible and portable way. It can be run locally, in a computer cluster, or in the cloud, improving the automation and the scalability of Bayesian modeling. We used Stan-NF to run our Bayesian regression models and get maximum a posteriori (MAP) estimates for *β*.

### Benchmark in simulated data

We benchmarked that approach on simulated data. We used the Flux simulator (Griebel et al. 2012) to generate simulated bulk RNAseq data. The Flux simulator first simulates a transcriptome, that is, the distribution of number of molecules per transcript, according to a modified Zipf’s Law. That was derived from experimental evidence indicating that most transcripts are lowly expressed, while there are very few transcripts expressed in high levels. Then, given that distribution, it simulates an RNAseq experiment and the corresponding data.

In order to simulate matched whole cell, nuclear, and cytosolic data, we first utilized the Flux simulator to assign randomized expression levels to the transcripts annotated in the human genome (GRCh38) by GENCODE (v29). We assumed these to be the “whole cell” transcript expression values. Subsequently, for each transcript, we distributed the number of molecules into “nuclear” and “cytosolic” fractions, such that the resulting nucleo-cytosolic distribution has a specific predetermined *β* parameter. From the same whole cell transcriptome, we generated different sets of nuclear and cytosolic transcriptomes, each time with an increasing bias for the cytosolic fraction (with values of *β* equal to 0.5, 0.6, 0.7, and 0.8 correspondingly). Then we performed simulated RNAseq experiments based on the simulated transcriptomes separately for the whole cell, the nucleus and the cytosol. Finally, we processed the resulting RNAseq dataset with the Grape-NF pipeline (https://github.com/guigolab/grape-nf) for quantifying transcript expression (see Methods).

We then applied our Bayesian regression approach to estimate *β*. The estimated values were essentially identical to the “real” (simulated) ones: 0.5, 0.59, 0.7, and 0.8, respectively. Then, we used the estimated *β* to calculate the localization index (eq. 8) for every transcript. The distribution of the localization index strongly resembles that of the simulated fraction of RNA molecules per transcript localized in the cytosol, independently of the overall nucleo-cytosolic distribution of the RNA volume (i.e. the *β* parameter, Fig. 2A).

**Figure 2.**
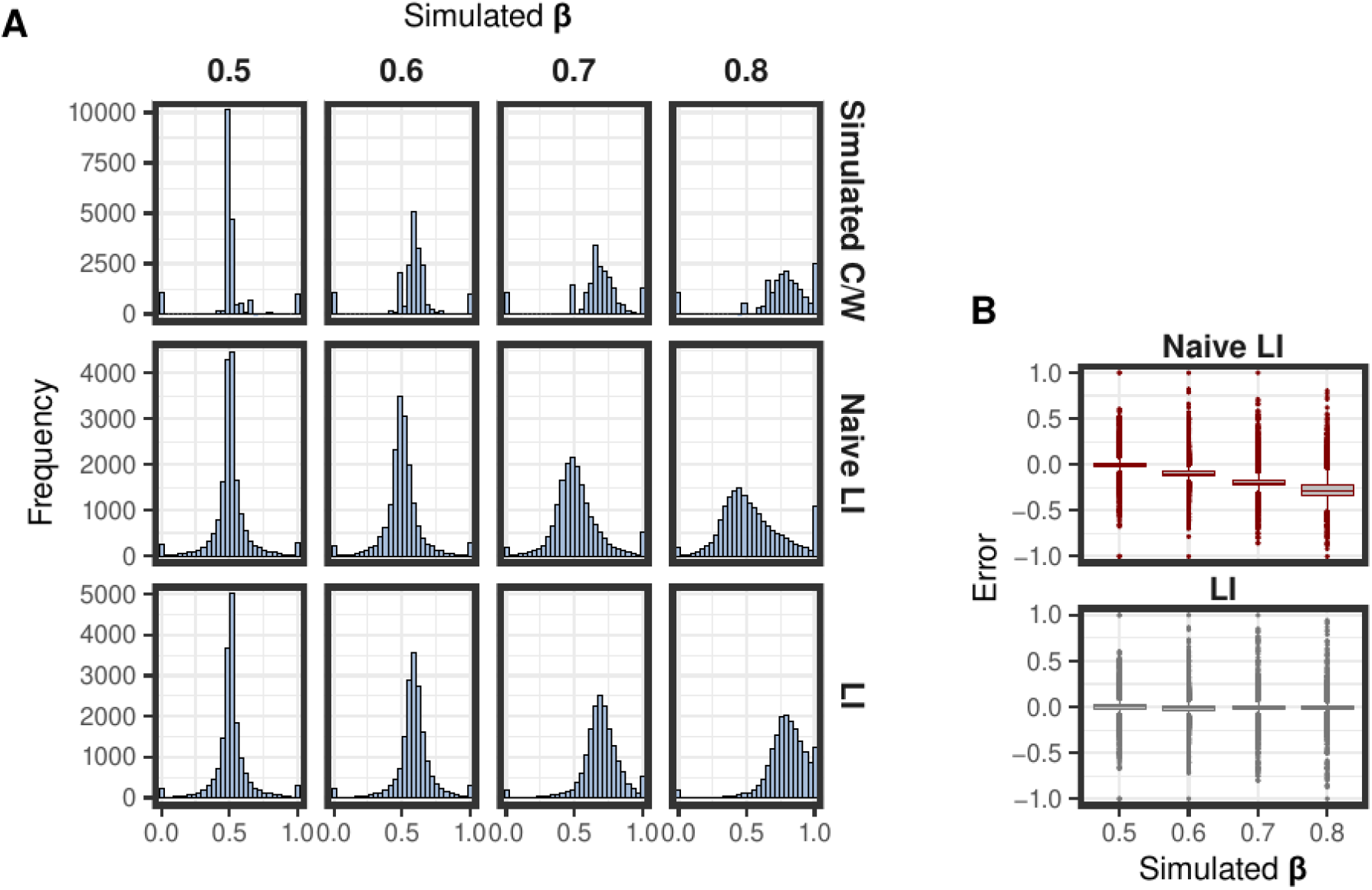
Localization index assessment on the simulated data. The simulation includes four samples with different nucleo-cytosolic RNA distribution (simulated *β* : 0.5 − 0.8). (A) In the upper row of panels there is the distribution of the proportion of the total RNA molecules, across the simulated transcriptome, localized in the cytosol. The histograms of the lower row of panels illustrate the distribution of the calculated localization index for the simulated transcripts, after estimation of the *β*. (B) Boxplots showing the error of estimating transcript localization using the localization index (left panel), or the naive localization index (right panel). The error is the deviation of the calculated index per transcript from the simulated proportion of cytosolic RNA molecules per transcript.

Additionally, we calculated the naive localization index based on the nuclear and cytosolic FPKMs as indicated in equation (9). When comparing this index with the simulated localization, we observed an increasing error, as the overall nucleo-cytosolic distribution deviated from the equilibrium (*β >* 0.5, Fig. 2B). In contrast, the error in the localization index (calculated using the estimated *β*) remained around zero on average (Fig. 2B).

### Benchmark in single cell RNAseq data

There are very few experimental systems with whole cell, nuclear and cytosolic RNAseq data been produced and the fraction of the cytosolic RNA volume (*β*) independently estimated. Recently, however, Abdelmoez et al. constructed a microfluidics system in order to isolate nuclear and cytosolic RNA from single cells, and perform RNAseq. They used their system to measure RNA expression in K562 cells. We took advantage of that dataset, which includes whole-cell, nuclear and corresponding cytosolic RNAseq from K562 single cells. For each fraction, we constructed a virtual bulk RNAseq sample by pooling the sequencing output from different cells, and processed that data to quantify transcript expression. We used our bayesian regression approach, and estimated *β* to be 0.82, very close to the average proportion of transcript abundance in the cytoplasm (0.84) reported by Abdelmoez et al. 2018.

### Analysis of the whole cell, nuclear and cytosolic RNAseq data produced by ENCODE

One of the richest publicly available datasets including whole cell, nuclear, and cytosolic RNAseq from the same biological samples has been produced as part of the ENCODE project (The ENCODE Project Consortium 2012), and it contains data from eleven different cell lines in biological replicates (Djebali et al. 2012). Analyses of these data can provide some insight on the preferential subcellular localization of different transcript types.

We used our Bayesian regression approach, and estimated *β* for each replicate available in every cell line (Fig. 3A). In most of these cell lines the estimated *β* is highly consistent between the two replicates, and almost always greater than 0.7. That indicates that in these cells a large fraction of the cellular RNA is in the cytosol. That is in accordance with previous studies (Perry et al. 1974; L Johnson et al. 1975; Herman et al. 1976; FUJIWARA et al. 2010; FUJITA et al. 2012) that measured the proportion of polyA+ RNA localized in the nucleus, with methods orthogonal to RNA sequencing, at 0.2-0.3. In six of the twenty one replicates available, we estimated *β >* 0.95 (the two replicates from A549, and one replicate each from GM12878, IMR-90, K562 and HeLa-S3). While these values are possible, we have eliminated these samples from further analysis as they could likely arise from a combination of limited efficiency in the experimental fractionation protocol and of instability of our Bayesian regression approach. We used *β* to calculate the localization index across transcripts in each of the remaining ten cell lines (Fig. 3B). For the cell lines with two replicates only transcripts present in both replicates were kept for further analysis, and the average of the localization index was calculated.

**Figure 3.**
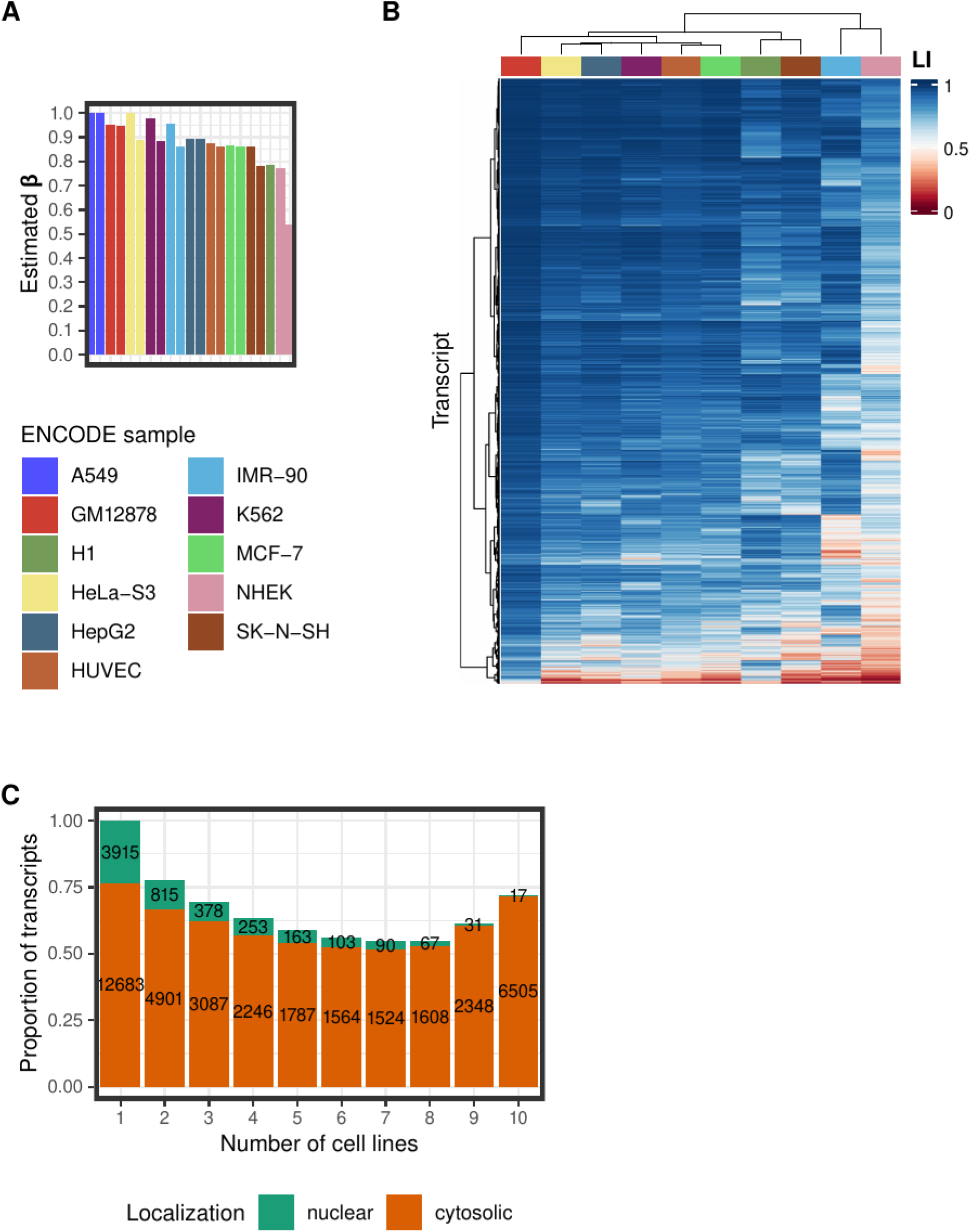
Transcript localization in the ENCODE dataset. (A) Estimated *β* values (i.e. the fraction of the total RNA volume that localizes to the cytosol) for every replicate of all bio-samples in the dataset. For further analysis, we kept the samples with estimated *β ≤* 0.95. (B) In the remaining 10 cell lines, the localization index (LI) for every transcript was calculated (averaged across replicates, when available). The heatmap depicts LI values for 9,074 transcripts that are expressed in all 10 cell lines. Hierarchical clustering with Ward’s method was performed to group the bio-samples and the transcripts. (C) The barplot depicts the proportion of the transcripts that have the same localization, nuclear (localization index (*LI*) *≤* 0.5) or cytosolic (*LI ≥* 0.5), across all the cell lines in which they were found to be expressed. The number of transcripts each proportion corresponds to is also indicated. The x-axis displays the number of cell lines that the corresponding number of transcripts were detected.

We first investigated to what extent the cellular localization of a transcript is consistent across biological conditions. For each number of cell lines in which a given transcript is expressed, we computed the number (and proportion) of transcripts which are always cytosolic (*LI ≥* 0.5) and which are always nuclear (*LI ≤* 0.5, Fig. 3C). For instance, we found 9,074 transcripts that are expressed in all ten cell lines. Of these 6,505 (72%) are cytosolic in all of them and 17 are nuclear in all of them (0.2%). The proportion of transcripts that are always nuclear decreases with the number of cell lines in which the transcript is expressed, as expected since the probability that, in an experiment, all binary trials have the same outcome decreases with the number of trials. This is not the case, however, for transcripts that are always cytosolic as, counterintuitively, the proportion of transcripts that are always cytosolic is the largest for transcripts that are expressed in all the ten cell lines. This strongly suggests that transcripts that are constitutively expressed tend to be systematically localized in the same cellular compartment, while transcripts that are transiently expressed or that are expressed in a more restricted manner have a less consistent cellular localization.

Next, we examined the distribution of the localization index for different transcript biotypes (Fig. 4A, pooled across cell types, Supplemental Fig. S1 for each cell type separately). As expected, transcripts from protein coding genes are more cytosolic than long non-coding RNAs whose distribution is more skewed towards the nuclear fraction. Transcripts annotated as “retained introns” have a strong nuclear distribution, suggesting that these transcripts are captured by RNAses before splicing is complete, or that intron retention could be a mechanism contributing to prevent translocation of transcripts from the nucleus to the cytosol (Chang and Sharp 1989; Legrain and Rosbash 1989; Kwiatek et al. 2023). Consistently with that, we also observed that processed pseudogene transcripts are significantly more cytosolic than unprocessed pseudogene transcripts (Supplemental Fig. S1).

**Figure 4.**
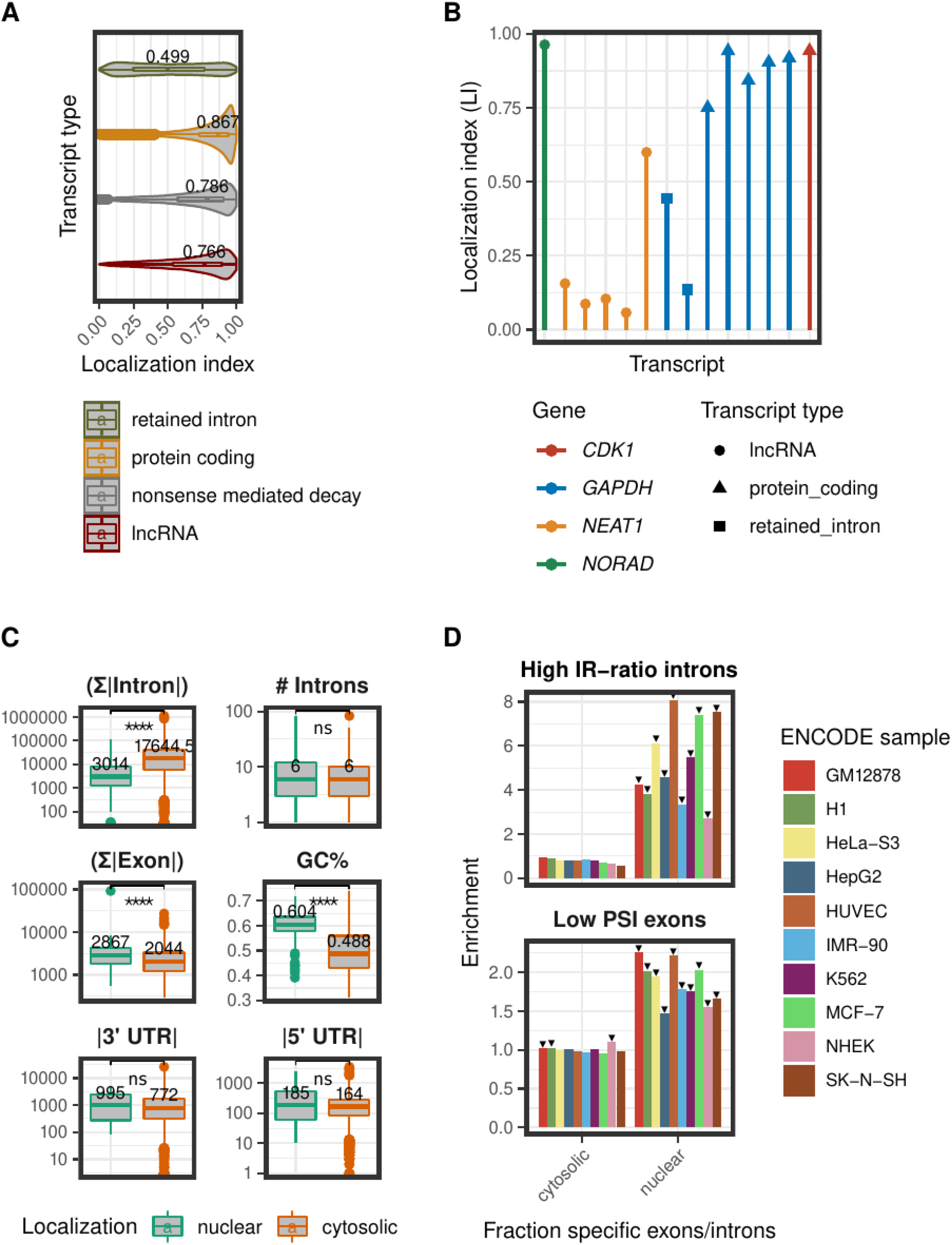
Association of the localization index (LI) with different transcript features. (A) The LI distribution for different transcript biotypes. Data shown for the 4 biotypes with the highest number of expressed transcripts. LI values were pooled across all the cell lines. The median LI value is indicated for every transcript biotype. (B) LI values for the expressed transcripts of four well-known genes in the SK-N-SH cell line. (C) Association of the LI with different features of transcript sequence and structure. The boxplots compare the total intron length, total number of introns, total exon length, GC content, 3’ UTR length, and 5’ UTR length, between nuclear (*LI <* 0.4) and cytosolic (*LI >* 0.6) transcripts. The transcripts compared had the same localization across all the samples, in which their expression was detected. The Wilcoxon rank sum test was used to assess the statistical significance of the observed differences. Significance levels: “ns” p-value *>* 0.05, **** p-value *≤* 0.0001. (D) Splicing efficiency in nuclear and cytosolic transcripts. The upper graph depicts the enrichment of high IR-ratio introns (*IR*-*ratio >* 0.5) in the set of localization specific introns for every cell line analyzed. The lower graph shows the enrichment of low PSI exons (*PSI <* 0.5) in the set of localization specific exons for every cell line analyzed. The statistical significance of the enrichment was assessed with a hypergeometric test followed by the Bonferroni correction. Statistically significant enrichments are indicated with a black arrow.

As an example, in Figure 4B we show the localization index calculated for the transcripts of four different genes in the SK-N-SH cell line. As it is possible to see, within the same gene we observe different nucleo-cytosolic localizations depending on the transcript biotype. For *GAPDH*, transcripts with retained introns are much more nuclear than protein coding transcripts. In other cases, different transcripts with the same biotypes from the same gene show different localization. For instance for *NEAT1*, a known nuclear lncRNA, there is one transcript with a more cytosolic distribution.

Thus, we investigated next how often transcripts from the same gene were preferentially located in different compartments. Within a given cell line, we considered a transcript cytosolic if its localization index was larger than 0.6 and nuclear if it was smaller than 0.4. The number of cytosolic transcripts ranged from 13,073 (57.6% of expressed transcripts in the cell line) to 27,517 (74.3%, Supplemental Table S1). The numbers of nuclear transcripts ranged from 252 (0.9%) to 5,430 (18%). For most cell lines, the percentage of genes with transcripts in different localizations ranged from 36% to 49%. When transcripts from the same gene localized in different compartments, they were most often of different biotypes. However, we also found genes with transcripts of the same biotype preferentially localized in different compartments. For most cell lines, the percentage of such genes ranged from 7% to 15% (Supplemental Table S1). This suggests that cellular localization is, to some extent, a property of the transcript rather than of the gene. That is also supported by the observation that the localization of transcripts annotated as “retained intron” is similar between “retained intron” transcripts of protein coding genes and “retained intron” transcripts of lncRNA genes (Supplemental Fig. S2).

We also identified a set of transcripts with very strong cellular localization. From the transcripts expressed in 5 cell lines or more, we selected those with very strong cytosolic localization (localization index *>* 0.9 in all cell lines where expressed, 345 transcripts) and those with very strong nuclear localization (localization index *<* 0.3 in all cell lines where expressed, 81 transcripts). Gene Ontology functional enrichment analysis on the strong cytosolic transcripts revealed that these transcripts are enriched for functions related to translation. This is reasonable, because the translational machinery is one of the most fundamental components of the cell, and especially for immortalized cell lines, such as those from ENCODE. While most of the strong cytosolic transcripts are protein coding (270, 78%), we also found 14 lncRNAs, and 23 transcripts annotated as “nonsense mediated decay”. Interestingly, many of those 14 lncRNAs were found in the literature to be associated with cytosolic localization and microRNA regulation (Han et al. 2019; Tang et al. 2020; YX Chen et al. 2021; Zheng et al. 2021; Rosano et al. 2022; Yu et al. 2023). Moreover, while most of the strongly nuclear transcripts are retained intron annotated transcripts (64 retained intron annotated, 2 lncRNA, 3 nonsense mediated decay), we found 10 protein coding transcripts that are strongly nuclear. All of them belong to genes that also have cytosolic transcripts.

Finally, we investigated whether features of transcript structure and sequence differ between the consistently cytosolic and consistently nuclear transcripts (Fig. 4C). We found that the total intronic length was larger in cytosolic than in nuclear transcripts. We observed the opposite trend for total exonic length. Cytosolic transcripts have also lower GC content than nuclear transcripts. We found small differences in UTR length, which however were not statistically significant.

Intron structure and splicing efficiency have been associated with RNA subcellular localization in previous studies (Zuckerman and Ulitsky 2019; Zeng and Hamada 2020). To investigate splicing efficiency in cytosolic and nuclear transcripts, we calculated the percent spliced in (PSI) score for all exons, and the intron retention ratio (IR-ratio) for all introns (see Methods). We assessed the enrichment of low PSI exons (PSI *<* 0.5), and high IR-ratio introns (IR-ratio *>* 0.5) in the cytosolic and nuclear fractions. To do this, we focused on the sets of exons and introns that are cytosolic-specific, and nuclear-specific. In other words, those within each cell line that are present only in cytosolic transcripts (*LI >* 0.6) or only in nuclear transcripts (*LI <* 0.4). We measured enrichment values by controlling for the global frequency of high IR-ratio introns (or low PSI exons). For instance, the enrichment of high IR-ratio introns in the set of nuclear-specific introns was calculated as follows:

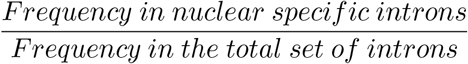

As illustrated in Figure 4D, introns with high IR-ratio were overrepresented in the set of nuclear-specific introns, and exons with low PSI were overrepresented in the set of nuclear-specific exons.

## Discussion

The strong compartmentalization is a characteristic feature of the eukaryotic compared to the prokaryotic cell. Among the subcellular compartments, the nucleus, which contains most of the cellular DNA, is arguably the main eukaryotic compartment. The nuclear membrane physically separates two fundamental biological processes: the transcription of DNA to RNA in the nucleus, and the translation of RNA to proteins in the cytosol. The need for mRNA molecules to be exported from the nucleus into the cytosol offers additional opportunities for gene regulation, as mRNA retention in the nucleus limits the rate of protein synthesis. To fully understand the relevance and the characteristics of this eukaryotic-specific regulatory layer, accurate measurements of the abundance of transcript species in the nucleus and in the cytosol are required.

Here we approach the problem of predicting the relative cytosolic (nuclear) abundance of transcripts by using RNA sequencing. We show that if we produce whole cell, nuclear and cytosolic RNAseq in a given sample, we can estimate the relative abundance of a given transcript in the nucleus and in the cytosol. We first employ Bayesian linear regression to estimate the RNA volume (i.e. the total number of nucleotides in RNA molecules) in the cytosol (nucleus). For most cell lines investigated here, we estimated that between 70% and 90% of the cellular PolyA+ RNA resides in the cytosol, a value which is consistent with values previously reported in the literature (Perry et al. 1974; L Johnson et al. 1975; Herman et al. 1976; FUJIWARA et al. 2010; FUJITA et al. 2012).

In a few cases, our estimates on cytosolic RNA are unrealistically high. We attributed this to a combination of factors. First, cell fractionation experiments enrich for nuclear and cytosolic RNA, but they are not able to completely segregate the two fractions, or they enrich them unevenly. Second, we estimate the relative volume of cytosolic RNA from tens of thousands of linear equations (each one relating the expression of a transcript in the nucleus and the cytosol with the whole cell expression). The intrinsic error in the determination of expression levels from RNAseq experiments (in particular for lowly expressed genes), may make the equations inconsistent, leading, on occasions, to unstable estimates. Nonetheless, the consistency of our estimates of cytosolic (nuclear) RNA across biological replicates argues that they are informative of the actual cellular volumes.

Next, we used the estimated relative cytosolic RNA volume to estimate the proportion of the total RNA molecules localized in the cytosol for each transcript independently (the localization index). Exhaustive benchmarks in simulated data support our estimates. With all the caveats discussed, we believe that these estimates constitute a unique resource to investigate the underlying molecular determinants of the relative cellular localization of transcripts genome-wide. Thus, we have found that the nucleo-cytosolic distribution of transcripts is partially encoded in their sequence, with nuclear transcripts significantly more GC rich and with shorter intronic regions compared to cytosolic transcripts. Thus, the cellular localization appears to be more a property of the transcripts than of the genes. Consistently, we have found that a large proportion of genes (up to 50% depending on the cell line) generate transcripts that are in different preferential localizations in the same cell line. While these transcripts tend to belong to different biotypes, we also found that many genes (up to 15% depending on the cell types) generate transcripts of the same biotype that are in different preferential localizations. These results argue for a complex model of genic functionality, in which the same gene might be involved, simultaneously, in a variety of biological functions, carried out by transcripts in different compartments.

Remarkably, against statistical expectations, we have found that transcripts expressed in many cell lines tend to have a constant cellular localization. That is, transcripts that are constitutively expressed tend to be localized in the same compartment (essentially the cytosol) across biological conditions, while transcripts expressed only in a few conditions tend to have a more variable localization. Transcripts constitutively expressed are likely to be associated with core cellular functions, less susceptible to regulation in a condition specific way than transcripts that are transiently expressed or expressed only in a few conditions. These transcripts may be subjected to many layers of regulation in a condition specific way, including regulation of transcript translocation to the cytosol.

In summary, we developed a method that estimates the transcript nucleo-cytosolic distribution. Importantly, this method can be easily applied to existing data. This would involve reassessing current findings, transitioning from a paradigm focused on compartment enriched transcripts to accurately determining the localization proportions for each transcript. We believe that our work constitutes, with all the caveats, one of the most accurate resources to date to investigate RNA localization and trafficking, a central problem in the regulation of gene expression.

## Methods

### Simulated data

We simulated whole cell, nuclear, and cytosolic bulk RNAseq data (Supplemental Fig. S3) using the flux simulator (Griebel et al. 2012, version 1.2.1). Flux simulator initially generates randomized transcript expression levels for transcripts annotated in a genome based on a modified Zipf’s Law, that was derived by fitting a model to experimental datasets. We first generated a simulated number of molecules per transcript for the whole cell sample.

We used the human genome annotation GENCODE v29 (Frankish et al. 2022) on the human genome assembly GRCh38, including only the canonical chromosomes from the nuclear genome in order to simulate a total of 5 million RNA molecules (on average 24.19 molecules per transcript, ranging from 0 to 38,112 molecules). Next, we split the number of molecules per transcript into nuclear and cytosolic fractions. The difference between the nuclear and cytosolic fractions for each transcript was sampled from a negative binomial distribution with specified mean and dispersion parameters, in order to control the total nucleo-cytosolic distribution. We generated four different splits, each one with a nucleo-cytosolic distribution characterized by a *β* parameter of 0.5, 0.6, 0.7, and 0.8 accordingly. In each of these splits, there are nuclear and cytosolic fractions with expression levels assigned to each transcript.

Subsequently, we applied the Flux simulator again to generate RNAseq reads originating from the transcripts in each fraction in each of the splits and in the original whole cell simulated transcriptome. Library preparation was simulated with the default values, which correspond to a uniform random RNA fragmentation, and a reverse transcription using random hexamers as primers. Sequencing was simulated with paired-end reads of a 76 bp length. For every sample the total amount of sequenced reads was roughly 50 million. Lastly, we analyzed the simulated RNAseq datasets using the Grape-NF pipeline (https://github.com/guigolab/grape-nf, version 1.1.1). The pipeline was run with the default parameters of the standard profile for mapping with STAR (Dobin et al. 2012), and transcript quantification with RSEM (Li and Dewey 2011). As a result, we produced FPKM values for the transcriptome in each fraction in each split, and for the whole cell transcriptome.

### Single cell RNAseq data

We fetched the single-cell RNAseq data from the Sequencing Read Archive (accession: SRP119800). This dataset includes single-cell RNAseq data generated from total (available for 12 cells), nuclear (43 cells), and cytosolic RNA (43 cells). Since the method applied here requires a correspondence between the whole cell RNAseq data and the fractionated RNAseq data, we used the nuclear and cytosolic RNAseq data for only 12 out of the 43 available cells. The identifiers of the files in the used dataset can be found in Supplemental Table S2. We utilized the fastp utility (S Chen et al. 2018, version 0.20.1) to preprocess and check the quality of the raw data. Then, we merged the sequencing data of all the cells for the whole-cell, nuclear, and cytosolic samples, in order to create a pseudo-bulk RNAseq dataset. Finally, we mapped the raw reads on the human genome (assembly: GRCh38) and estimated transcript quantification (transcript annotation: GENCODE v24) for each pseudo-bulk RNAseq sample using the Grape-NF pipeline, similarly to how it was done for the simulated data described above.

### ENCODE data

The dataset analyzed (Djebali et al. 2012) was generated by the group of Thomas Gingeras as part of the ENCODE project. It consists of polyA selected bulk RNA sequencing data (library fragment size *>* 200 bp, paired-end reads) from 9 different cell lines (A549, GM12878, H1, HeLa-S3, HepG2, IMR-90, K562, MCF-7, SK-N-SH) and 2 different primary cell samples (HUVEC, NHEK). For every sample, there are available whole cell, cytosolic, and nuclear RNAseq data at the ENCODE portal (Luo et al. 2019). We downloaded transcript quantification information, as generated by the STAR-RSEM (Li and Dewey 2011; Dobin et al. 2012) pipeline (Genome assembly: GRCh38, Genome annotation: GENCODE v29). The identifiers of the files in the used dataset can be found in Supplemental Table S3.

### Data preprocessing

For every sample in the simulated, pseudo-bulk, and ENCODE datasets, we utilized the transcript expected count, and the effective length values from the RSEM output. First, we assessed the reproducibility of transcript quantifications between replicates using non-parametric irreproducible discovery rate (see Djebali et al. 2012, npIDR). For each replicate, we kept transcripts with a npIDR value lower than 0.1 in all three fractions (whole cell, nuclear, cytosolic). This step was skipped for the H1 cell line data, which include only one biological replicate, as well as for the simulated and pseudo-bulk datasets. In addition, we calculated counts per million (CPM) for every transcript, in order to filter out transcripts with considerably low expression. For every replicate, we kept transcripts whose expression in the whole-cell and at least one of the two fractions, cytosol or nucleus, was greater than or equal to 1 CPM. Furthermore, we filtered out transcripts encoded in the mitochondrial DNA, as these should not affect the distribution of the RNA between nucleus and cytosol. Finally, fragments per kilobase of transcript per million fragments mapped (FPKM) values were calculated per transcript.

For downstream analyses of the ENCODE dataset, we eliminated six out of the twenty one replicates available, that had estimated *β >* 0.95 (the two replicates from A549, and one replicate each from GM12878, IMR-90, K562 and HeLa-S3).

### Splicing analysis

We used the RNAseq data from the whole-cell fraction in order to calculate the percent spliced in (PSI) score and the intron retention ratio (IR-ratio). We fetched the mapped reads (bam files) for every sample in the whole-cell dataset using the ENCODE portal (accession numbers of the used files can be found in Supplemental Table S3). PSI estimates the occurrence of exon skipping events. To measure PSI for all exons, we utilized IPSA-NF (https://github.com/guigolab/ipsa-nf, version 4.2). IPSA-NF is a Nextflow (Tommaso et al. 2017) implementation for the Integrative Pipeline for Splicing Analyses, which operates within a framework that extends the PSI metric to encompass diverse categories of splicing events (Pervouchine et al. 2012). IR-ratio estimates the occurrence of intron retention events. To measure IR-ratio for all introns, we utilized IRFinder (Lorenzi et al. 2021, version 2.0.0) in BAM mode. IRFinder calculates IR-ratio as intronic abundance divided by the sum of intronic abundance and normal splicing abundance.

### Estimation of *β*

We took advantage of the relationship described in equation (7), so as to estimate *β* using a regression approach. Regression was performed under a Bayesian framework as follows:

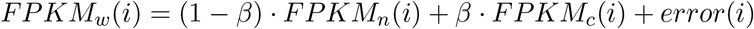

where the *FPKM* values are the measured transcript quantifications in the whole cell (*w*), nuclear (*n*), and cytosolic (*c*) fractions. We modeled the error term as Student-t distributed, in order to make our estimations more robust:

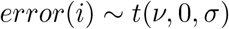

Moreover, we chose the following priors for the model parameters:

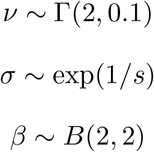

where *ν* is the degrees of freedom, *σ* is the error scale, *s* is the standard deviation of the *FPKM*_*w*_ across transcripts, and *β* is the regression coefficient. The prior for the degrees of freedom was proposed by Juárez and Steel 2010. The prior for the error scale *σ*is a weakly informative data dependent prior, an approach implemented also in the rstanarm R package (Goodrich et al. 2022). The prior for the regression coefficient *β* is a Beta distribution, which is symmetric, centered around 0.5 and bounded between 0 and 1. The chosen distribution implies equally distributed RNA between cytosol and nucleus.

Bayesian linear regression was implemented using the Stan probabilistic programming language (Stan Development Team 2021). Specifically, we used CmdStan (version 2.28.0), a command-line interface to Stan. We designed Stan-NF (https://github.com/vntasis/stan-nf), a reproducible and portable pipeline around CmdStan, using Nextflow workflow manager (Tommaso et al. 2017). CmdStan can be used to compile models written in Stan, perform Markov chain Monte Carlo (MCMC) sampling from the posterior distribution, calculate diagnostics for every MCMC run, and generate summaries of the results. Stan-NF was designed in order to automate, and scale all that. It can compile multiple Stan models in parallel, and then run multiple MCMC chains on multiple data input for every model in parallel.

Stan-NF was run using the no-U-turn sampler (NUTS), a variant of the Hamiltonian Monte Carlo (HMC) algorithm, in order to sample from the posterior distribution of *β*. We used 4 chains and obtained 6,000 samples per chain, out of which the first 2,000 were used for warm-up. Lastly, we specified the value of 0.9 for the target average proposal acceptance probability (adapt_delta), and 10 for the maximum treedepth (max_depth). The final point estimate for *β* was the maximum a posteriori (MAP) estimate.

### Estimation of the localization index

We defined the localization index for any transcript *x* as the portion of the total amount of RNA molecules of that transcript in the cell that are localized in the cytosol. It was estimated based on the formula of the equation (8):

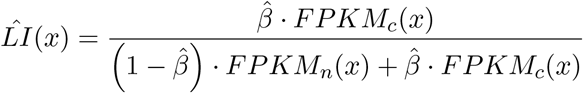

where 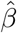 is the *β* parameter estimated by our regression approach, and *FPKM* values are the measured transcript quantifications.

### Transcript type annotation

We used the GENCODE v45 transcirpt annotation to retrieve transcript biotype labels, as it is improved compared to the GENCODE release 29 with which the data was initially processed. The transcripts with no correspondence between the two annotation versions were ignored for the association of the transcript biotype with the localization of a transcript (Fig. 4A).

### Gene Ontology functional enrichment

For functional enrichment analysis we focused on the groups of strongly cytosolic and strongly nuclear transcripts. That is, transcripts that are expressed in 5 cell lines or more, have consistent localization in all cell lines, with *LI >* 0.9 (cytosolic) or *LI <* 0.3 (nuclear). For the assessment of the enrichment of Gene Ontology terms in the set of strongly localized transcripts, we used g:Profiler (Kolberg et al. 2023). As the background set of transcripts (statistical domain scope), we used all the transcripts expressed in 5 cell lines or more. Lastly, we assessed statistical significance with a g:SCS threshold of 0.05.

### Calculation of transcript features

We used BEDTools (Quinlan and Hall 2010, version 2.31.0) and Python GTF Toolkit (Lopez et al. 2019, version 1.6.2) in order to compute the following transcript features from the genome (annotation: GENCODE v29, assembly: GRCh38): total intron length, total exon length, total number of introns, length of the untranslated regions, GC content.

### Splicing and localization

We studied splicing efficiency of the localization-specific exons and introns. For every cell line, we selected the exons and introns that were found only in nuclear transcripts (*LI <* 0.4), or only in cytosolic transcripts (*LI >* 0.6). We assessed the enrichment of low PSI exons (PSI *<* 0.5), and high IR-ratio introns (IR-ratio *>* 0.5) within the groups of localization-specific exons and introns. To do this, we also measured the frequency of low PSI exons, and high IR-ratio introns among all detected exons and introns, to use them as the expected frequencies. We calculated the enrichments as follows:

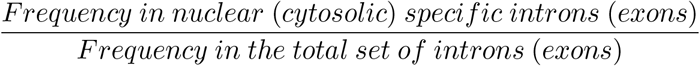

## Code Availability

For the bigger part of the analysis, we used the R programming language (R Core Team 2021, version 4.0.4). The code necessary to regenerate the analysis, and the results of this study can be found in the following repository: https://github.com/vntasis/localization_index_paper. Moreover, we used containerization software (Docker (Merkel 2014) and Singularity (Kurtzer, Sochat, et al. 2017; Kurtzer, cclerget, et al. 2020)) in order to make the analysis reproducible. The images with the software required to run the code are accessible (details can be found in the code repository).

## Acknowledgements

This work and its publication were supported by the National Institute of Health - NIH under grant agreement n° 2 U24 HG007234-09 and the Agència de Gestió d’ajuts Universitaris I de Recerca (AGAUR) under grant agreement n° 2021-SGR2021-01304. V.F.N. is supported by the Ministerio de Ciencia, Innovación y Universidades de España under grant agreement PRE2019-088504. We acknowledge the support of the Spanish Ministry of Science and Innovation to the EMBL partnership, Centro de Excelencia Severo Ochoa and CERCA Programme / Generalitat de Catalunya. We thank R. Garrido and R. Carbonell Garcia (CRG) for administrative support.

